# Multicolor Light-Induced Immune Activation via Polymer Photocaged Cytokines

**DOI:** 10.1101/2022.10.03.510638

**Authors:** Lacey A Birnbaum, Emily C. Sullivan, Priscilla Do, Biaggio Uricoli, Christopher C Porter, Curtis J Henry, Erik C Dreaden

**Affiliations:** Coulter Department of Biomedical Engineering, Georgia Institute of Technology and Emory University; Molecular and Systems Pharmacology Graduate Program, Emory University School of Medicine; Department of Hematology and Medical Oncology, Winship Cancer Institute of Emory University; Department of Pediatrics, Emory School of Medicine; Aflac Cancer and Blood Disorders Center of Children’s Healthcare of Atlanta; Petit Institute for Bioengineering and Bioscience, Georgia Institute of Technology

**Keywords:** cytokines, bioconjugation, PEGylation, immunotherapy

## Abstract

Cytokines act as potent, extracellular signals of the human immune system and can elicit striking treatment responses in patients with autoimmune disease, tissue damage, and cancer. Yet despite their therapeutic potential, recombinant cytokine-mediated immune responses remain difficult to control as their administration is often systemic whereas their intended sites of action are localized. To address the challenge of spatially and temporally constraining cytokine signals, we recently devised a strategy whereby recombinant cytokines are reversibly inactivated via chemical modification with photo-labile polymers that respond to visible LED light. Extending this approach to enable both *in vivo* and multicolor immune activation, here we describe a strategy whereby cytokines appended with heptamethine cyanine-polyethylene glycol are selectively re-activated *ex vivo* using tissue-penetrating near-infrared (NIR) light. We show that NIR LED light illumination of caged, pro-inflammatory cytokines restores cognate receptor signaling and potentiates the activity of T cell-engager cancer immunotherapies *ex vivo*. Using combinations of visible- and NIR-responsive cytokines, we further demonstrate multi-wavelength optical control of T cell cytolysis *ex vivo*, as well as the ability to perform Boolean logic using multicolored light and orthogonally photocaged cytokine pairs as inputs, and T cell activity as outputs. Together, this work demonstrates a novel approach to control extracellular immune cell signals using light, a strategy that in the future may improve our understanding of and ability to treat cancer and other diseases.

## INTRODUCTION

Cytokines orchestrate a range of biological processes and act as key signaling molecules of the human immune system that regulate inflammation as well as immune cell migration, proliferation, and differentiation.^1-4^ Yet despite their central importance to immune homeostasis, the clinical approval and use of recombinant cytokine immunotherapies to-date has been hampered in part due to the pleiotropic effects of these molecules and their relatively low cell- or tissue-specificity that leads to dose-limiting or off-target side effects.^5^ For example, cardiopulmonary toxicities were observed in early clinical studies of aldesleukin (rIL-2) immunotherapy in cancer and poor treatment outcomes were associated with the drug-induced expansion of immunosuppressive Tregs.^6^ Likewise, early trials of rIL-12 and rIL-15 immunotherapy were limited by significant hematologic and hepatic toxicity, as well as macrophage activation syndrome and intense cytokine secretion, respectively.^7, 8^

Given their potent induction of antitumor immunity, various approaches to achieve cytokine cell- or tissue-specificity have been employed to-date and include cytokine secretion by engineered cells, as well as affinity targeting, polymer conjugation, protein mutation, and *de novo* protein design.^9, 10^ The use of various stimuli to locally activate cytokine prodrugs represents another promising approach to mitigate the off-target effects of cytokines and include approaches to activate cytokine fusion proteins^11-13^ and drug carriers^14, 15^ via proteolysis as well as those that release cytokines from protein-crosslinked nanoparticles in response to reducing conditions at the T cell surface.^16, 17^

To address the challenge of improving the cell- and tissue-specificity of recombinant cytokines, we recently developed a strategy whereby cytokine activity (e.g. IL-2, IL-12, IL-15) can be blunted via conjugation with a photo-labile polyethylene glycol (PEG) and recovered via brief, in this case blue, LED light exposure.^18^ While these studies demonstrated the feasibility of *in vivo* cytokine photoactivation using tissue phantoms, we concluded that the clinical applications of this approach were limited to cutaneous or subdermal tissues and those accessible by *in situ* light sources (e.g. catheter-guided fiber lasers) as a result of efficient absorption and scattering of light by tissues and biological fluids at visible wavelengths. In contrast to visible light, near-infrared (NIR) photos can more efficiently penetrate human tissues including bone, skin, and muscle up to several cm^19^ and diagnostic methods relying on NIR light sources have demonstrated clinical utility in optical coherence tomography (OCT) and near-infrared spectroscopy (NIRS) applications.

Leveraging the ability of NIR photons to efficiently penetrate human tissues, here we advance our prior approach^18^ to reversibly photocage immunostimulatory cytokines by using photo-labile PEG that de-shields in response to NIR rather than visible light. We show that LED light exposure of NIR-photocaged cytokines can be used to reversibly modulate cognate cytokine receptor signaling and enhance the activity of bispecific cancer immunotherapies that rely on T cell-dependent cytolysis. Using combinations of visible and NIR light-responsive cytokines, we further demonstrate orthogonal, multicolor photocontrol of T cell cytolytic activity *ex vivo* and the novel ability to perform Boolean logic using multicolor light and differentially photocaged cytokines. This work expands the working toolbox for protein photocaging and may be extended in the future to a range of other biological systems that rely on extracellular signaling or to synthetic cytokine/receptor pairs.

## RESULTS

To achieve NIR control of cytokine signaling, we first devised a heterobifunctional linker which would undergo photolysis to liberate one of two conjugation sites upon light exposure. Gorka *et al*. previously showed that C4’-dialkylamine substituted heptamethine cyanine fluorophores undergo regioselective photooxidation followed by C4’-N hydrolysis and cyclization to liberate aromatic alcohols following NIR light exposure.^20^ We surmised (i) that substitution of this leaving group with a phenolic tetrazine would enable the photo-induced loss of a variety of compounds appended via trans-cyclooctene (TCO) click chemistry and (ii) that addition of a distal azido ethoxy group to the backbone cyclohexene would enable the linker to be protein-anchored via orthogonal azide-alkyne click chemistry.^21-23^ As anticipated, the subsequent heterobifunctional linker (hereafter, **cCy**) exhibited strong NIR absorption and fluorescence (ex/em, 705/810 nm, **Figure S1,2**, as well as rapid photolysis as characterized via UV-Vis absorption spectroscopy following NIR irradiation (785 nm, photooxidation *t*_1/2_ 25 min; **Figure 1a,b, S3)**. To characterize the spatial resolution in which cCy can be photocleaved, we next immobilized it to TCO-functionalized glass slides via click chemistry and exposed these slides to collimated NIR LED light (730 nm) under aqueous conditions through a custom lithographic mask **(Figure 1c)**. Using the intrinsic fluorescence of cCy, we observed a lateral resolution for photocleavage ∼51 μm via epifluorescence imaging **(Figure 1c-e)**, a finding that is significant in that this dimension approaches that of single mammalian cells, thus demonstrating its potential utility in intravital imaging studies,^24-26^ as well as *ex vivo* or *in vitro* culture studies in the future.^27, 28^

**Figure 1.**
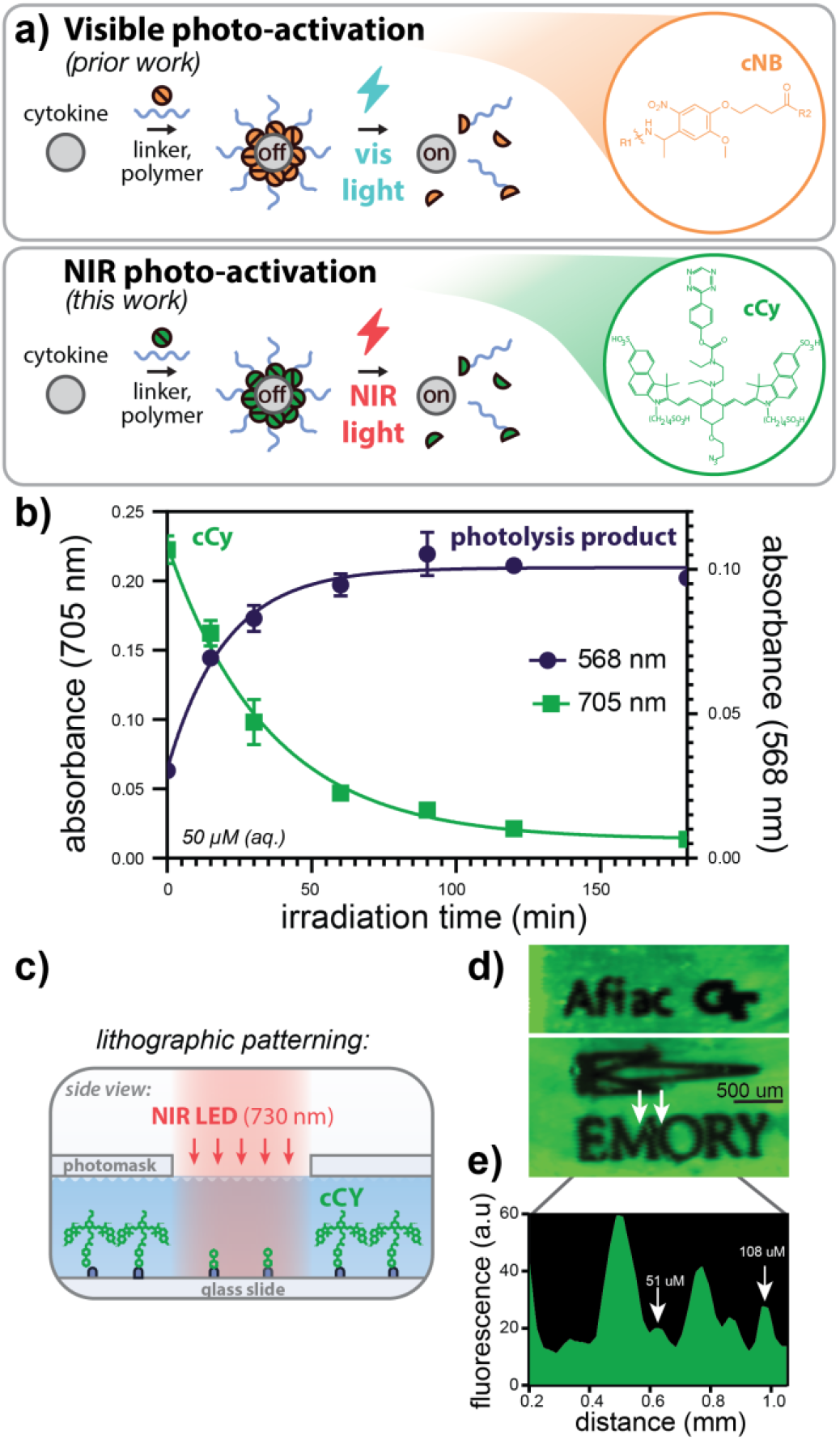
Near-infrared (NIR) photo-labile linker, cCy, undergoes rapid, spatially well-resolved photocleavage. **a)** Strategy for blue and NIR photon-gated protein activation via cCy-linked polymer conjugation. **b)** Aqueous NIR photolysis kinetics of cCy as measured by UV-Vis spectroscopy of parent (blue) and decomposition (purple) products during irradiation (785 nm, 110 mW/cm^2^). **c)** Schematic illustrating cCy immobilization to TCO-functionalized glass slides and subsequent aqueous-phase photolithographic patterning (730 nm, 100 mW/cm^2^).**d)** Fluorescence micrographs of cCy (green) photopatterns and **(e)** corresponding image line scans demonsrating approximately 51 μm lateral resolution (785 nm ex, 812-832 nm em). Data in (b) represent mean ± SD of three technical replicates per time point.

Having demonstrated that cCy undergoes rapid and spatially well-defined photolysis, we next investigated whether the conjugation of proinflammatory cytokines with cCy-polyethylene glycol (cCy-PEG) modulates their activity on T cells. Using carbodiimide and TCO-tetrazine chemistry, we modified surface primary amines of recombinant IL-12 with cCy or cCy-PEG_20kDa_ and, using reporter cells that secrete alkaline phosphatase in a pSTAT4-dependent manner, we identified cCy conjugation densities that increase the apparent molecular weight of rIL-12 whilst preserving its signaling activity **(Figure 2a-c)**. Subsequent NIR LED light exposure of IL-12-cCy led to a near complete restoration of IL-12 electrophoretic mobility and, using this optimized cCy density, we next conjugated IL-12 with cCy-PEG, observing a 95±6 -fold decrease in potency relative to rIL-12 as measured by pSTAT4 reporter cell activity **(Figure 2c,d)**. Following NIR photolysis, we observed a 66±7 -fold recovery in IL-12 potency (EC_50, 24h_). To confirm the generalizability of this approach to photocaging, we demonstrated a comparable increase and recovery in rIL-15 electrophoretic mobility following conjugation with cCy-PEG and NIR light exposure (**Figure S4**). Together, these data demonstrate that cCy-PEG reversibly modulates the size and *ex vivo* activity of rIL-12 on T cells in a NIR photon-dependent manner.

**Figure 2.**
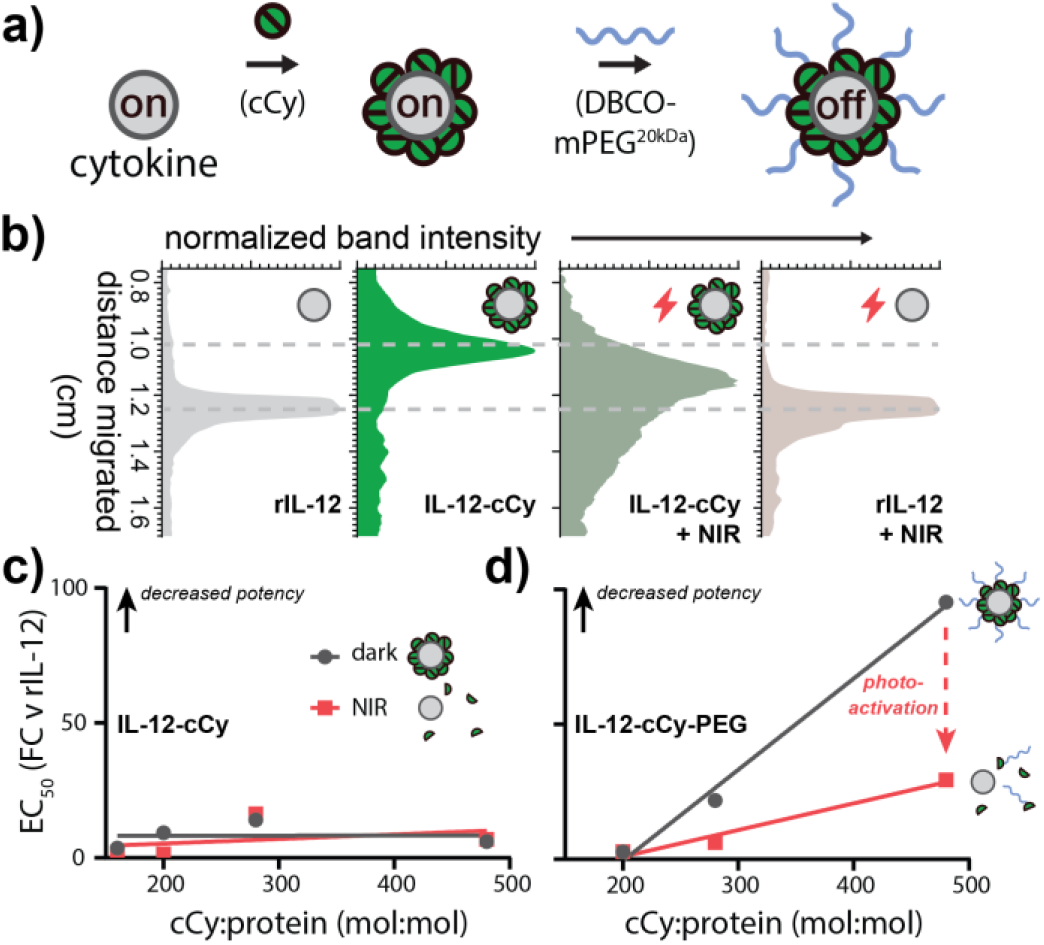
NIR light exposure of IL-12-cCy-PEG restores protein size and de-represses pSTAT4 signaling. **a)** Schematic illustrating the modification of recombinant IL-12 with cCy-PEG_20kDa_. **b)** Densitometry of IL-12 and IL-12-cCy with and without NIR light exposure illustrating a decrease then recovery in electrophoretic mobility following cCy conjugation and subsequent NIR light exposure, respectively (730 nm, 100 mW/cm^2^). IL-12 dose-dependent STAT4 activation with varying **(c)** cCy and **(d)** cCy-PEG conjugation ratio as measured via SEAP/chromogenic assay using STAT4 reporter cells (24 h) with and without NIR light exposure (730 nm, 100 mW/cm^2^) and normalized as fold-change (FC) relative to rIL-12. Data in (c,d) represent mean ± SD of three technical replicates.

In prior work, we demonstrated that recombinant IL-12 can enhance the cytolytic activity of the T cell bispecific immunotherapy, blinatumomab, currently approved to treat patients with relapsed and refractory B cell acute lymphoblastic leukemia (B-ALL).^29, 30^ To determine whether IL-12-cCy-PEG can likewise improve blinatumomab-induced cytolysis in an NIR light-dependent manner, we treated cocultures of labeled primary human CD8+ T cells and CD19+ NALM-6 B-ALL cells with blinatumomab and rIL-12 or IL-12-cCy-PEG, with and without NIR light exposure and measured CD19-specific cell lysis and T cell activation via flow cytometry and IFNγ ELISA, respectively **(Figure 3a-c)**. As anticipated, we observed no significant change in blinatumomab-induced cytolysis or T cell activation following co-culture with IL-12-cCy-PEG without illumination; however, following NIR LED exposure of the protein, we observed an increase in both CD19-specific lysis and IFNγ secretion that was statistically indistinguishable from that enhanced by rIL-12.

**Figure 3.**
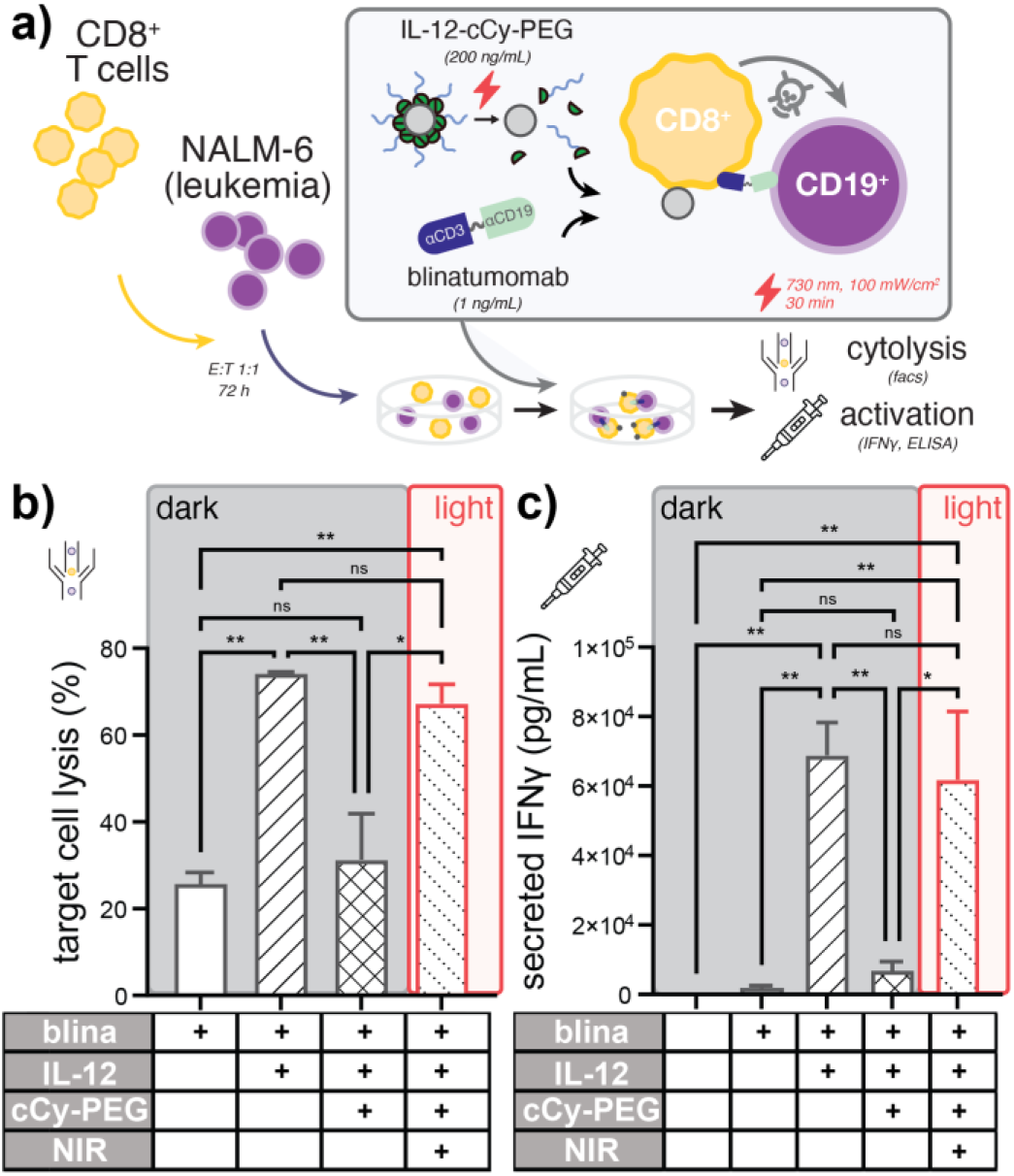
NIR light modulates the potency of blinatumomab immunotherapy when co-treated with IL-12-cCy-PEG. **a)** Schematic illustrating assay conditions in which co-cultures of labeled primary human CD8+ T cells and labeled CD19+ NALM-6 B-ALL cells are treated with blinatumomab and rIL-12 or IL-12-cCy-PEG, with and without NIR light illumination. **b)** CD19-specific target cell lysis demonstrating loss and recovery of blinatumomab cytolytic activity following cCy-PEG photocaging and subsequent NIR LED illumination (730 nm), respectively, as measured by flow cytometry. **c)** IFNγ measurements indicating a repression and de-repression of IL-12 dependent T cell activation following polymer photocaging and NIR light exposure, respectively, as measured by ELISA of co-culture supernatants. Data in (b,c) represent mean ± SD of at least two technical replicates from a representative healthy T cell donor.

After demonstrating that NIR photocaging can be used to modulate the influence of IL-12 on blinatumomab-induced cytolysis, we asked whether other cytokines might cooperate with IL-12 to further improve drug activity. Prior studies demonstrated (i) that IL-2 treatment upregulates IL-12R on T cells (ii) that IL-12 treatment upregulates IL-2R (CD25) on CD8+ T cells, and (iii) that the combination of both IL-12 and IL-2 synergize to improve tumor control in syngenetic mouse models of renal cell carcinoma.^31-33^ To determine whether cooperativity between IL-12 and IL-2 can further enhance blinatumomab activity *ex vivo*, we co-cultured CFSE-labeled primary human CD8+ T cells and CellTrace Violet-labeled CD19+ NALM-6 B-ALL cells with blinatumomab in the presence or absence of each recombinant cytokine or their combination **(Figure 4a)**. We observed that rIL-2 improved drug-induced cytolysis to levels comparable with rIL-12 alone and that both proteins additively combined with blinatumomab to augment CD19-specific cell lysis as measured by flow cytometry **(Figure 4b)**, despite that neither blinatumomab nor cytokine treatment impacted target CD19 or PD-L1 immune checkpoint ligand expression on leukemic B cells **(Figure S5)**. Interestingly, while rIL-12 and rIL-2 combined only sub-additively to alter T cell proliferation, they synergized to enhance IFNγ secretion as measured by dye dilution and ELISA of coculture supernatants, respectively **(Figure 4c,d)**. Together, these data support that (i) recombinant IL-2 alone may be used to enhance the potency of blinatumomab immunotherapy and (ii) that cytokine-enhanced T cell activation rather than T cell expansion is a dominant contributor to the combined impact of IL-12 and IL-2 on blinatumomab activity observed here.

**Figure 4.**
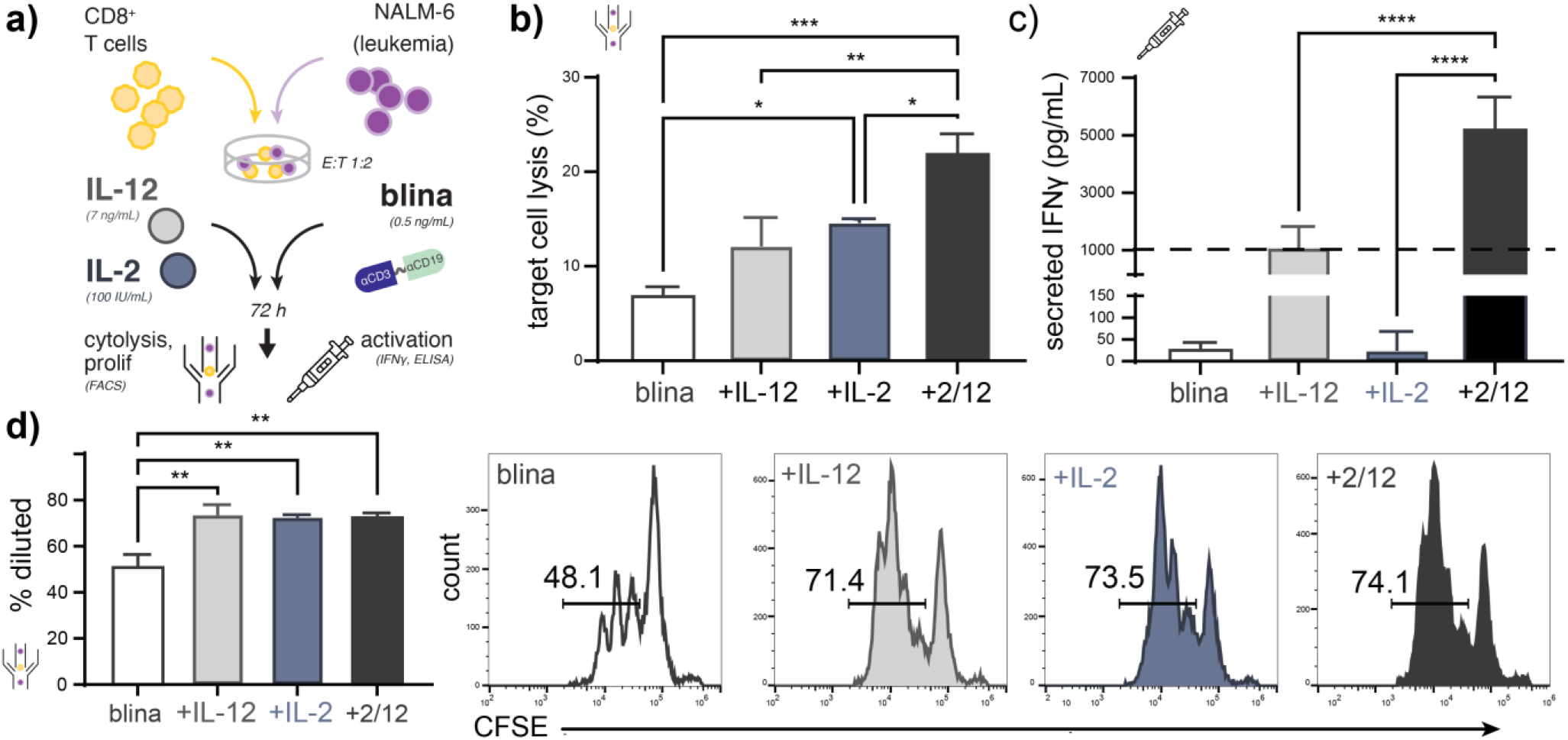
Recombinant IL-12 and IL-2 coopertively enhance blinatumomab-induced cytolytic activity and CD8 T cell activation *ex vivo*. **a)** Schematic illustrating assay conditions in which co-cultures of labeled primary human CD8+ T cells and labeled CD19+ NALM-6 B-ALL cells are treated with blinatumomab and rIL-12 or rIL-2. **b)** Blinatumomab-induced target cell lysis illustrating improved cytolytic activity from IL-12 or IL-2 cotreatment, as well as additive cooperative effects from their combination as measured by flow cytometry. Parallel measurements of **(c)** IFNγ production and **(d)** T cell proliferation demonsrating that rIL-12 and rIL-2 synergize to enhance CD8+ T cell activation but only sub-additively impact T cell proliferation as measured by ELISA of co-culture supernatants and flow cytometric dye dilution, respectively. The dashed line in (c) denotes the additive expectation as assessed via the Response Additivity model. Data in (b-d) represent mean ± SD of at least two technical replicates from a representative healthy T cell donor.

Having demonstrated that rIL-12 and rIL-2 cooperate to improve blinatumomab-induced cytolysis, we next asked whether the conjugation of each protein with wavelength-orthogonal photocages could be used to control their combined effect on CD8+ T cells, blinatumomab, and CD19+ leukemic B cells via multicolor LED light. In prior work, we developed a strategy to photocage IL-2 with blue LED light-responsive photocages based upon *o*-nitrobenzyl-linked PEG_20kDa_ (i.e. IL-2-cNB-PEG).^18^ Using this conjugate and the NIR-responsive IL-12 conjugate described here (IL-12-cCy-PEG), we constructed Boolean circuit that combines AND gates in which we digitize the presence or absence of NIR or visible LED light, as well as the presence or absence of discreet concentrations of IL-12-cCy-PEG or IL-2-cNB-PEG **(Figure 5a, S6)**. Using these components as inputs and blinatumomab-induced cytolysis level as a digitized output, we observed full concordance with the expected AND-AND gate truth table whereby above-threshold B cell lysis was observed only in the presence of both caged cytokines, as well as both visible and NIR LED light **(Figure 5b)**. Together, these data demonstrate that multicolor light can be used to interface with and modulate *ex vivo* immune responses when combined with photocaged cytokines.

**Figure 5.**
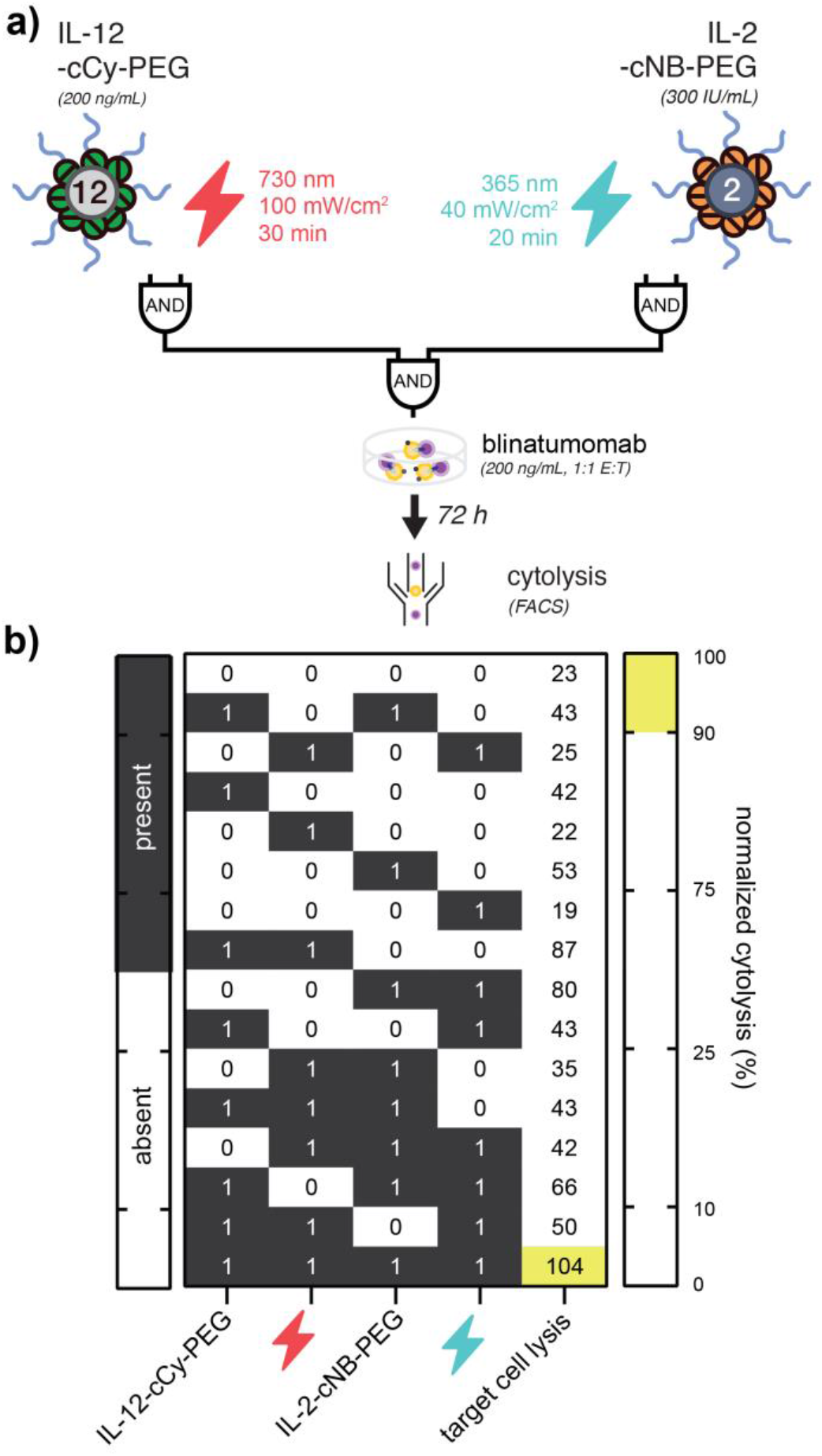
A combined photonic and biologic circuit based upon orthogonally photocaged IL-12 and IL-2. **a)** Schematic illustrating an AND-AND logic gate whereby (i) blue and NIR LED light, as well as IL-2-cNB-PEG and IL-12-cCy-PEG represent digitized inputs and (ii) normalized blinatumomab-induced target cell lysis respresents a digitized output. **b)** Truth table respresenting all possible combinations of initial states (green) and true outputs (blue), overlaid with observed output values of normalized drug-induced cytolysis as measured via flow cytometry. Data in (b) represent mean of 2-4 technical replicates from two healthy T cell donors.

## DISCUSSION AND CONCLUSIONS

Extending our prior approach to optically modulate cytokine signaling networks using visible LED light, here we describe a method (i) to photocontrol T cell cytokine signaling via tissue penetrant NIR light and (ii) to orthogonally modulate cooperative cytokine signaling via combined visible and NIR LED light. Using a photo-labile, hetero-bifunctional linker based upon heptamethine cyanine fluorophores (cCy), we demonstrated rapid NIR photolysis kinetics with a photooxidation half-life of ∼25 min, notably fast as compared with proteolytic activation methods that can exceed several hours in activation half-life. Using this novel photo-labile linker, we further observed a minimum lateral resolution for NIR photoactivation of ∼51 μm and anticipate further enhancements in resolution using focusing optics and/or multi-photon laser excitation (*e*.*g*. 1560 nm pulsed laser irradiation) as is common in high resolution microscopy applications.^34^ Such approaches when utilized with the NIR photocaging strategy describe here may find strong utility in intravital imaging applications^24-26^ as well as *ex vivo* studies of organoids and embryo-like body development where high spatial control of protein delivery is necessary for proper cell organization.^27, 28^

In prior work,^30, 35^ we demonstrated that rIL-12 could potentiate the therapeutic effects of the T cell engager, blinatumomab (Blincyto^®^) *ex vivo* and here we further demonstrate the NIR photo-control of this enhanced activity via NIR illumination of cCy-PEG conjugated rIL-12. This finding is significant as conventional rIL-12 therapies are well-known to be associated with severe hematologic and hepatic toxicities that may potentially compound with those observed from blinatumomab when delivered in combination. Using the approach described here, we observed a >98% reduction in signaling EC_50_ that we anticipate would mitigate such adverse effects *in vivo* via sustained or localized (*e*.*g*. bone marrow) delivery via NIR activation.

The observation that rIL-2, like rIL-12, potentiates the therapeutic activity of blinatumomab is further significant as this is the first report to our knowledge which support this effect from IL-2, as well as the first to demonstrate cooperativity between these two cytokines in enhancing the cytolytic activity of T cell bispecifics. Like IL-12, there are numerous IL-2-based clinical-stage drug candidates including protein fusions, PEG-cytokine conjugates, and mRNA.^9, 36, 37^ At the same time, there are now (4) clinically approved T cell bispecific immunotherapies that include both hematologic cancers and solid tumors.^38, 39^ Thus, these findings may support the future clinical development of combined, or chimeric, T cell bispecific and cytokine caner immunotherapies in the future.

Lastly, to exploit cooperativity between IL-12 and IL-2 in enhancing the *ex vivo* activity of blinatumomab, we constructed a Boolean circuit in which only 1 of 16 possible initial states – comprising caged IL-12, caged IL-2, blue light, or NIR light – would correspond to above-threshold cytolytic activity. In the future, this approach may be used to orchestrate or tune complex cytokine signaling networks using cocktails of caged cytokines and time- and/or wavelength-modulated light sources.

The finding that soluble proteins such as cytokines may be manipulated using exogenous light is also significant considering that the overwhelming majority of optical tools available to synthetic biologists such as photon-gated ion channels^40^ and light-dependent protein dimerization^41^ necessitate the use of live cells whose presence may be limiting to some systems or applications.

As discussed above, one salient barrier to the clinical application of recombinant cytokine immunotherapies in cancer is their relatively small size (typically 12-70 kDa) and therefore rapid excretion that necessitates frequent and/or high dosing and complex treatment management.^42, 43^ In prior work, the addition of just a single PEG molecule to GM-CSF or interferon alfa-2b was found to be sufficient to confer prolonged circulation and tissue-drug exposure that in-turn improves therapeutic index.^44^ Here, we likewise anticipate that prolonged circulation from IL-12-cCy-PEG relative to rIL-12 may enable it to more effectively elicit antitumor immune responses both alone and in combination with T cell bispecifics, immune checkpoint inhibitors, chimeric antigen receptor (CAR) T cells, and autologous gamma delta T cell^45^ or NK cell^46, 47^ therapies in the future where it may potentiate improved treatment outcomes.

In summary, here we demonstrate abiologic control of cytokine signaling networks using a multicolor light and a prodrug strategy whereby recombinant cytokines are chemically modified with photo-labile polymers that de-repress their activity in response to low-power LED light. Given recent advancements in the development of highly efficient and wavelength-variable photocages, ^48-50^ we anticipate further extensions of this approach in the future that will expand wavelength multiplexing capabilities and further exploit the utility of high spatial and temporal control over immune cell activation, migration, and differentiation.

## SUPPORTING INFORMATION

cCy chemical structure, absorption/emission spectrum, UPLC-MS, NMR analysis, and photolysis kinetics. Treatment-induced surface protein expression. IL-15 conjugation with cCy-PEG and NIR photo-cleavage.

## ACKNOWLEDGEMENTS

This work was supported in part by the American Cancer Society (#IRG-17-181-04), the Winship Cancer Institute, the National Institutes of Health Research Training Program in Immunoengineering (T32EB021962), the AAI Careers in Immunology Fellowship Program, the Coulter Department of Biomedical Engineering, and the Aflac Cancer and Blood Disorders Center of Children’s Healthcare of Atlanta. We are also grateful for assistance from the Children’s Healthcare of Atlanta and Emory University’s Pediatric General Equipment & Specimen Processing Core and the Pediatrics and Winship Flow Cytometry Core, as well as for helpful discussions with M. Quadir (NDSU).The content here is solely the responsibility of the authors and does not necessarily represent the official views of the organizations acknowledged herein.

## AUTHOR CONTRIBUTIONS

L.A.B., E.C.S., P.D., C.C.P., C.J.H., and E.C.D. designed research; L.A.B., E.C.S., P.D., B.U. and E.C.D. performed research or analyzed data; and L.A.B., C.C.P., C.J.H., and E.C.D. wrote the manuscript.

## COMPETING INTERESTS

L.A.B., P.D., and E.C.D. are inventors on a patent application related to this work describing the use of photolysis to activate therapeutic protein.

## METHODS

### Materials and Supplies

Unless otherwise specified, reagents were used as received without further purification. Recombinant human IL-2 (200-02, Peprotech), recombinant human IL-15 (570308, Biolegend), recombinant mouse scIL-12 (130-096, Miltenyi Biotech) recombinant human IL-12 (CT050-HNAH, Sino Biological). Sulfo-Cyanine7 NHS ester (Lumiprobe), DBCO-NHS ester (1160, Click Chemistry Tools), NHS-PEG-TCO ester (A137-10, Click Chemistry Tools), poly(ethylene glycol) methyl ether DBCO (20 kDa, A120-100, Click Chemistry Tools), poly(ethylene glycol) methyl ether azide (20 kDa, Nanocs). Polyacrylamide gels (Bio-Rad, 4-16 wt%). IFNγ ELISA (430104, Biolegend). CD19 antibody (BD Biosciences Clone SJ25C1) and PD-L1 antibody (eBioscience Clone MIH1).

### Cell lines and Primary Cells

HEK-Blue IL-12 cells (Invivogen) were cultured according to manufacturer’s recommendations in DMEM (Corning) supplemented with 10% heat inactivated FBS (VWR) and 100 IU/mL pen-strep (VWR). NALM-6 cells (gifted from Dr. Lia Gore, University of Colorado) were cultured in RPMI 1640 supplemented with 10% FBS and 100 U/mL penicillin and 100 μg/mL streptomycin. Primary CD8+ T cells were obtained from normal donor human buffy coats (LifeSouth) via Ficoll-Paque (Cytavia) gradient selection for peripheral blood mono-nuclear cells followed by human CD8+ T cell negative magnetic selection (Stem Cell Technologies). Primary CD8+ T cells were thawed in supplemented RPMI 1640 2-12 hours prior to the start of experiments.

### cCy Synthesis

2-((E)-2-((E)-2-((2-(((4-(1,2,4,5-tetrazin-3-yl)phenoxy)carbonyl)(ethyl)amino)ethyl)(ethyl)amino)-5-(2-azidoethoxy)-3-((E)-2-(1,1-dimethyl-7-sulfo-3-(4-sulfobutyl)-1,3-dihydro-2H-benzo[e]indol-2-ylidene)ethylidene)cyclohex-1-en-1-yl)vinyl)-1,1-dimethyl-7-sulfo-3-(4-sulfobutyl)-1H-benzo[e]indol-3-ium, cCy, was prepared at-scale by eMolecules (San Diego, CA) from 1,4-dioxaspiro[4.5]decan-8-one and purified to >96% via preparatory HPLC. cCy was structurally characterized by ^1^H NMR, ^13^C NMR, and UPLC-MS using a Acquity BEH C-18 column (1.7 μm): buffer A, 5mM ammonium acetate in water; buffer B, acetonitrile; gradient was 10-90 %A over 6 minutes at 0.3 mL/min flow rate. *See Supplementary Information for additional details*.

### cCy Spectroscopic Characterization

cCy was reconstituted to 50 mM in dry DMSO (Thermo Fisher Scientific) and further diluted 1000x in 1X PBS (Corning) for characterization studies. Absorption and emission spectra were measured on a Spectramax iD3 plate reader (Molecular Devices) at 10 nm increments. Photolysis kinetics were monitored using a NanoDrop One spectrophotometer following irradiation with a 785 nm diode laser (Crysta, 110 mW/cm^2^). Data was fit using a one phase decay model in Graphpad Prism.

### Photolithographic Patterning

1 mm silicone spacers with 9 mm diameter circular openings (Electron Microscopy Sciences) were adhered to amine coated glass slides (Nanocs). The slides were functionalized with NHS-TCO at a 10:1 NHS:amine ratio overnight at RT. Wells were then washed 6x with deionized water, 3x with 0.2% SDS, and again 3x with deionized water followed by conjugation with cCy at a 3:1 cCy:amine ratio overnight at RT in TBST. Wells were then washed 5x with TBST (0.1% Tween20) and covered in 50% glycerol. A chrome-coated quartz photomask was then placed on the silicone isolator and a collimated 730 nm LED (Thor Labs) was used to irradiate the slide for 30 minutes (100 mW/cm^2^). The irradiated slide was then washed 5x with TBST and mounted with Prolong Diamond antifade media (Life Technologies) for imaging. Patterned slides were imaged on an Odyssey CLX fluorescence imaging scanner. Image features were quantified and normalized using ImageJ.

### Conjugation with cCy-PEG

Cytokines were modified with cCy-PEG using carbodiimide and TCO-tetrazine chemistry, respectively using methods adapted from Perdue et al.^18^ Briefly, rIL-12 was sequentially reacted overnight with a 10-120-fold molar excess of NHS-TCO followed by a 40-480-fold molar excess of cCy and a 160-1,920-fold molar excess of DBCO-mPEG_20kDa_ (4°C with 800 rpm rotatory agitation). Excess reactants were removed via 7 kDa size exclusion chromatography (Zeba Spin, Thermo Fisher Scientific) in some experiments. IL-15-cCy-PEG was prepared using methods identical to IL-12-cCy-PEG except for the use of a 10-20-fold molar excess of NHS-TCO, 40-80-fold molar excess of cCy, and 160-320-fold molar excess DBCO-mPEG_20kDa_ (*see Supplementary Information*).

### Conjugation with cNB-PEG

rIL-2 was modified with cNB-PEG using carbodiimide and azide-alkyne chemistry, respectively using methods described in Perdue et al.^18^ Briefly, rIL-2 was diluted in 150 mM sodium phosphate buffer (pH 8.5) containing 0.5 mM SDS and sequentially reacted with 80-fold molar excess of cNB followed by N_3_-mPEG_20kDa_ at a 400-fold molar excess in PBS (4°C with 800 rpm rotatory agitation, overnight).

### Photocaged Cytokine Characterization

Protein electrophoretic mobility was characterized via polyacrylamide gel electrophoresis (4-16% weight) under reducing conditions. Protein bands were stained with Coomassie G250 (Bio-rad) and visualized using a Licor CLx imager. Coomassie was imaged in the 700 channel of the imager while the cCy was imaged in the 800 channel.

### Cytokine Photoactivation

Irradiation of IL-12-cCy-PEG was performed in transparent film-covered U-bottom 96 well plates following illumination with a 730 nm LED (ThorLabs, 100 mW/cm^2^, 30 min). cCy-PEG conjugates were diluted to 0.125-0.15 µg/mL in HEPES buffered saline (Invitrogen, pH 7.4) containing 0.1% BSA, 0.02% Tween 20 prior to irradiation. Irradiation of IL-2-cNB-PEG was similarly performed except for the use of a 365 nm LED (ThorLabs, 40 mW/cm^2^, 20 min) and pre-dilution in to 0.15 µg/mL in PBS (pH 7.4). Illumination was performed in all cases at 4 °C.

### Cytokine Receptor Signaling Assay

HEK-293 SEAP reporter cells were plated at 5×10^4^ cells per well in complete growth media within 96-well plates or at 2.5×10^4^ cells per well in a 384 well plate and treated with equimolar amounts of rh or caged cytokine, with and without light exposure. After 24 hours, 20 μL of supernatant was withdrawn per well, diluted 10-fold in Quanti-Blue reagent (rep-qbs, Invivogen), and placed at 37 °C for 0.25-3.0 hours before reading absorbance at 620 nm (Spectramax iD3 plate reader). Dose response curves were calculated using Graphpad Prism with a 4-parameter fit and normalized to background and signal from unmodified proteins.

### T Cell Cytolysis Assays

CD19+ NALM-6 cells (B cell acute lymphoblastic leukemia) were stained with Cell Trace Violet (Thermo Fisher Scientific) and primary human CD8+ T cells were stained with either Cell Trace Yellow (Thermo Fisher Scientific) or CFSE (Tonbo Biosciences) for ≥1 hours prior to co-culture. During co-culture, CD8+ T cells and 5×10^4^ NALM-6 cells were plated with 0.5-1 ng/mL blinatumomab (Invivogen) and cytokine, if applicable, in a U-bottom 96 well plate for 72 hours (1:1 or 1:2 E:T, as indicated). Cytokine concentrations ranged from 5-1000 IU/mL (IL-2), 7-400 ng/mL (IL-12). After 72 hours, cells were stained with Near IR Live/Dead stain (Thermo Fisher Scientific) for viability and immediately assessed by flow cytometry. Specific lysis was calculated as %Specific Lysis= 100*[(TreatedCTV+LD+)-(UntreatedCTV+LD+)]/(100-UntreatedCTV+LD+) where LD is viability stain, CTV is Cell Trace violet, and Untreated condition received no blinatumomab. Supernatant from these cultures was immediately frozen at -80° C. For IFNγ ELISA (Biolegend), co-culture supernatants were diluted 10-100-fold and assayed according to manufacturer’s instructions. During logic gate experiments, positive output was based upon inter-donor variability and defined as cytolysis within 50 %CI of values obtained from combined treatment with rIL-12, rIL-2, and blinatumomab. To assess changes in tumor cell phenotype, cells were stained for PD-L1 and CD19 for 1 hour at a 1:100 antibody dilution, washed, and immediately assessed by flow cytometry.

### Statistical Analysis

Unless otherwise noted, all p-values were calculated using either one-way ANOVA with Tukey post-hoc correction or one-way T test (Graphpad Prism), depending on the number of experimental conditions.

