## Supplementary Information for "Multicolor Light-Induced Immune Activation via Polymer Photocaged Cytokines"

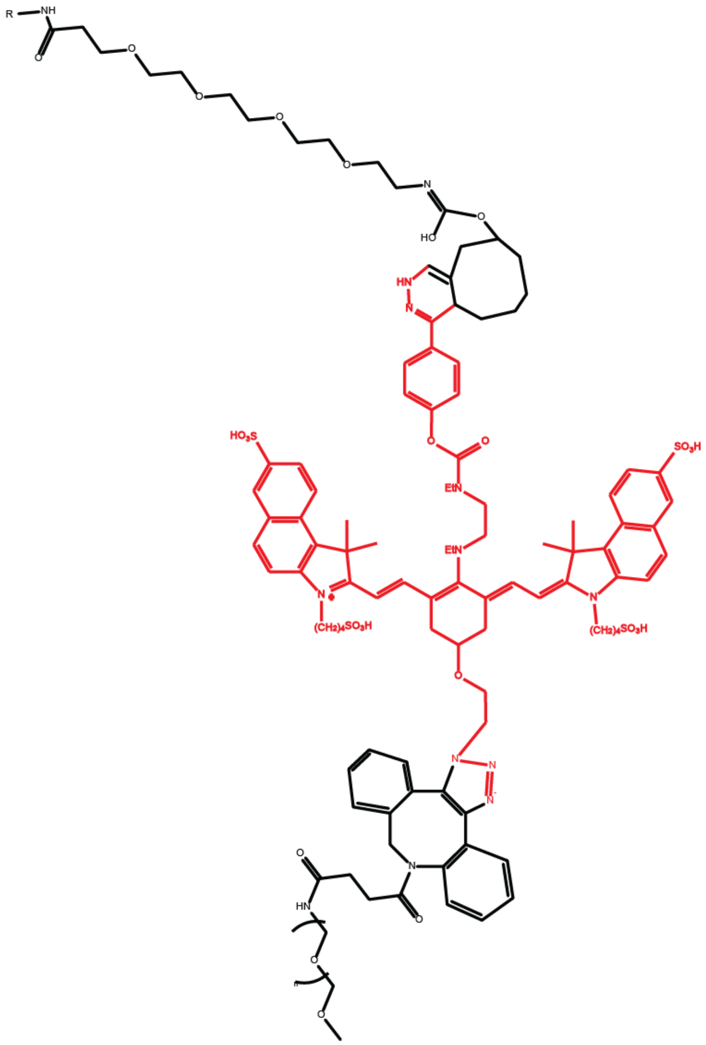

**Figure S1.** Structure of cCy-PEG following conjugation to cytokine surface amines.

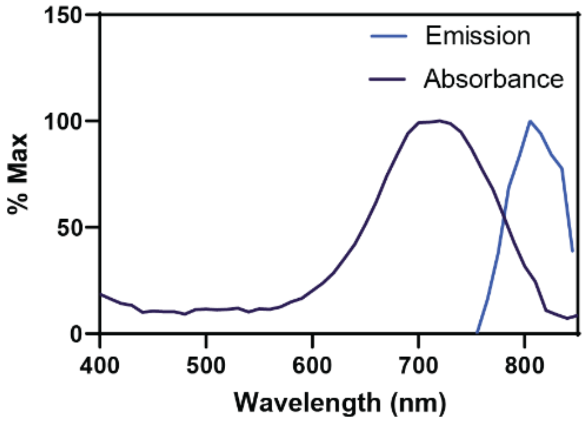


**Figure S2.** cCy absorption spectrum and emission spectra. Max emission is 810 nm (50µM, aq).

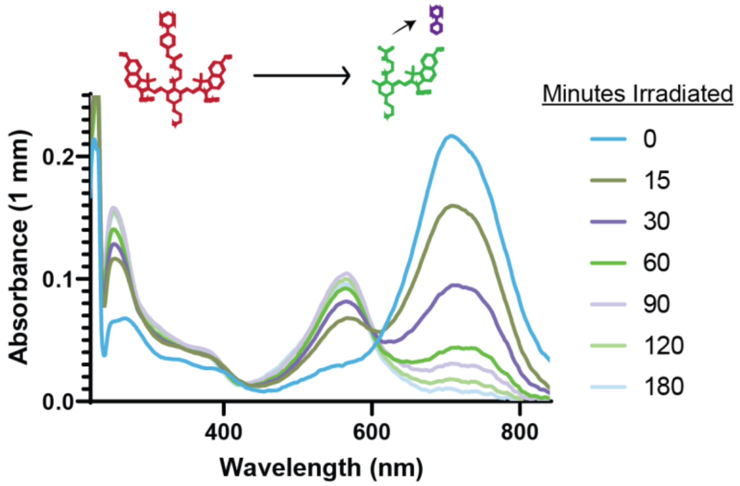


**Figure S3.** Spectral evolution of cCy UV-Vis-NIR absorption following NIR photolysis (785 nm, 110 mW/cm$^2$).

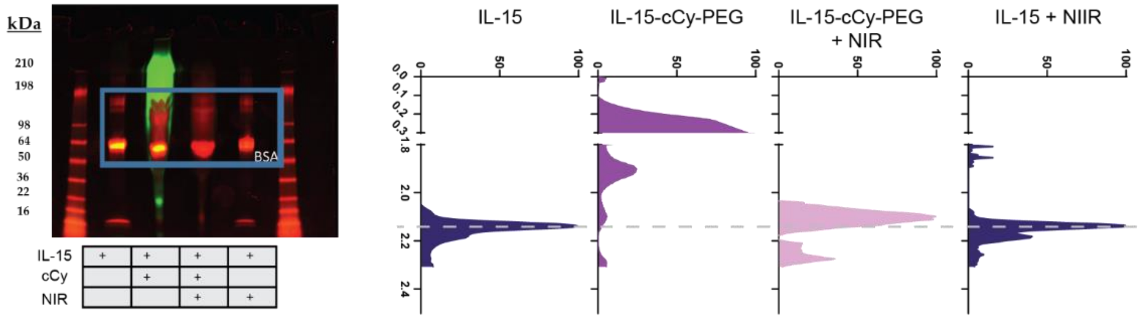


**Figure S4.** Conjugation of rIL-15 with cCy-PEG illustrating a decrease then recovery in electrophoretic mobility following cCy-PEG conjugation and subsequent NIR light exposure, respectively (730 nm, 100 mW/cm$^2$). *Left:* SDS-PAGE gel with neat protein in red (Coomassie) and to cCy in green (Cy7 channel). *Right:* Densitometry of rIL-15 and IL-15-cCy-PEG with and without NIR light exposure. BSA, added post-conjugation for stabilization, was omitted from densitometry plots for clarity.

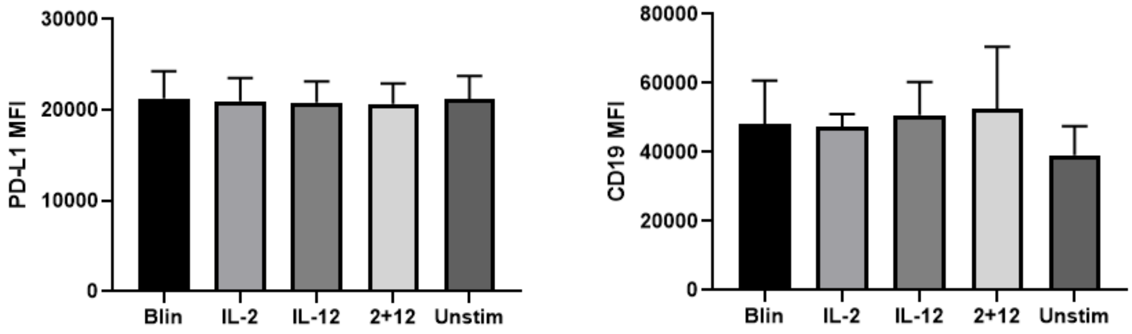


**Figure S5.** CD19 or PD-L1 expression on leukemic B cells is not significantly altered by blinatumomab or cytokine treatmnet. MFI of PD-L1 (left) and CD19 (right) expression on Nalm-6 leukemic B-cells following 1:1 E:T co-culture with CD8+ T cells with or without blinatumomab or cytokines (300 IU/mL IL-2 and 200 ng/mL IL-12) as-indicated. Data represent mean±SD of 6 technical replicates from 2 health donors.

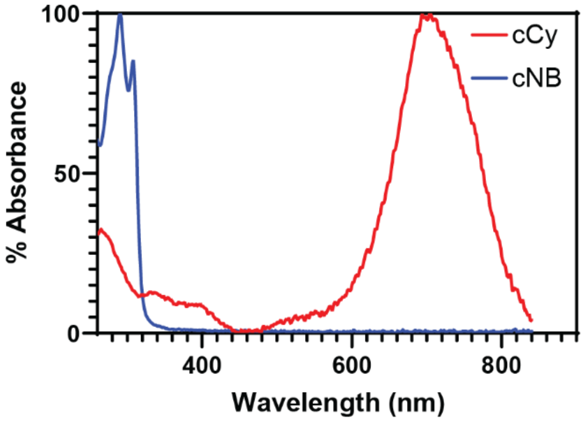


**Figure S6.** Optical absorption spectra for cCy (heptamethine cyanine, this work) as compared to cNB (*o*-nitrobenzyl, prior work). 50 μM (aq.) and 66.66μM (aq.), respectively.

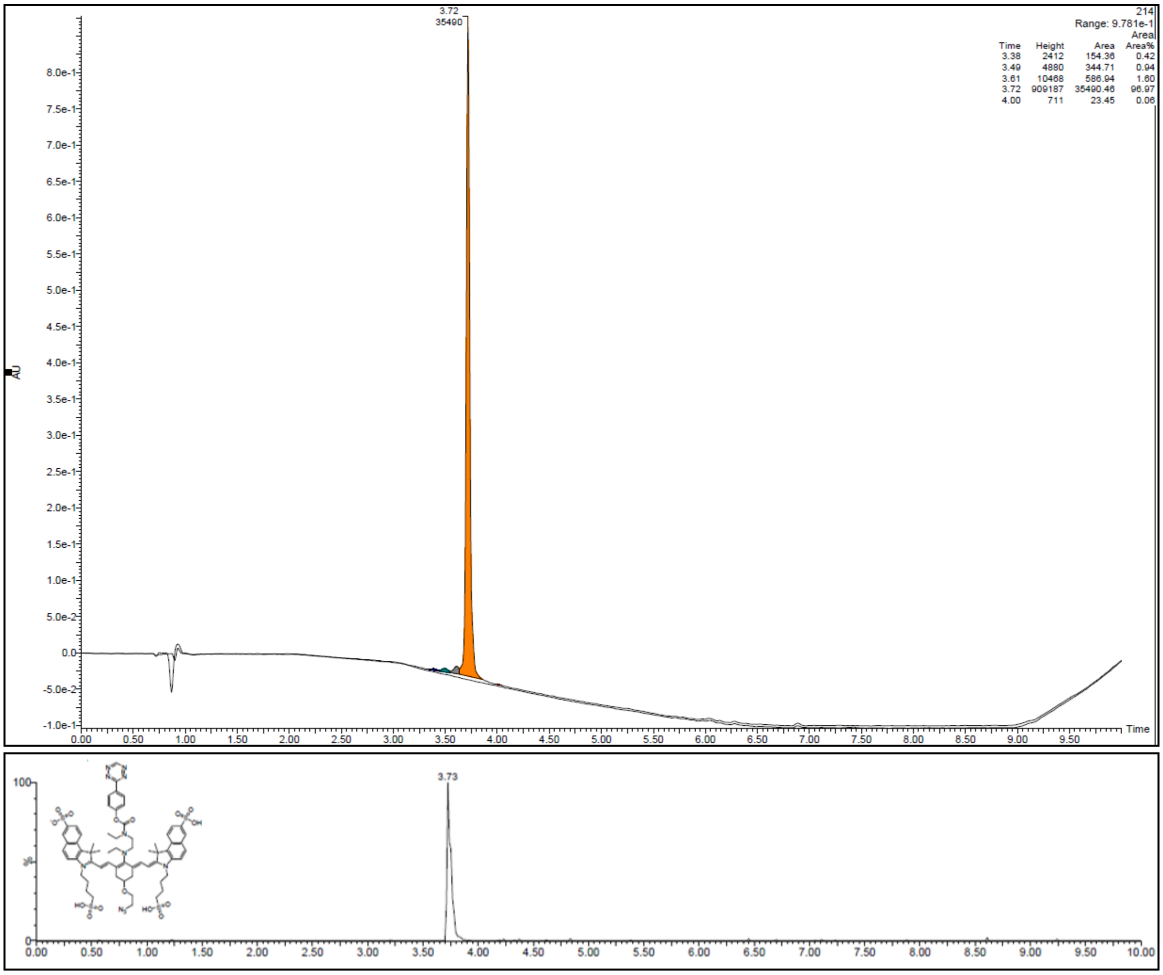


**Figure S7.** Top: LC-MS chromatogram of purified cCy by UV detector monitored at 214 nm. Peak purity 96.97%. Bottom: Chromatogram of elution of 1352.558 molecular weight peak.

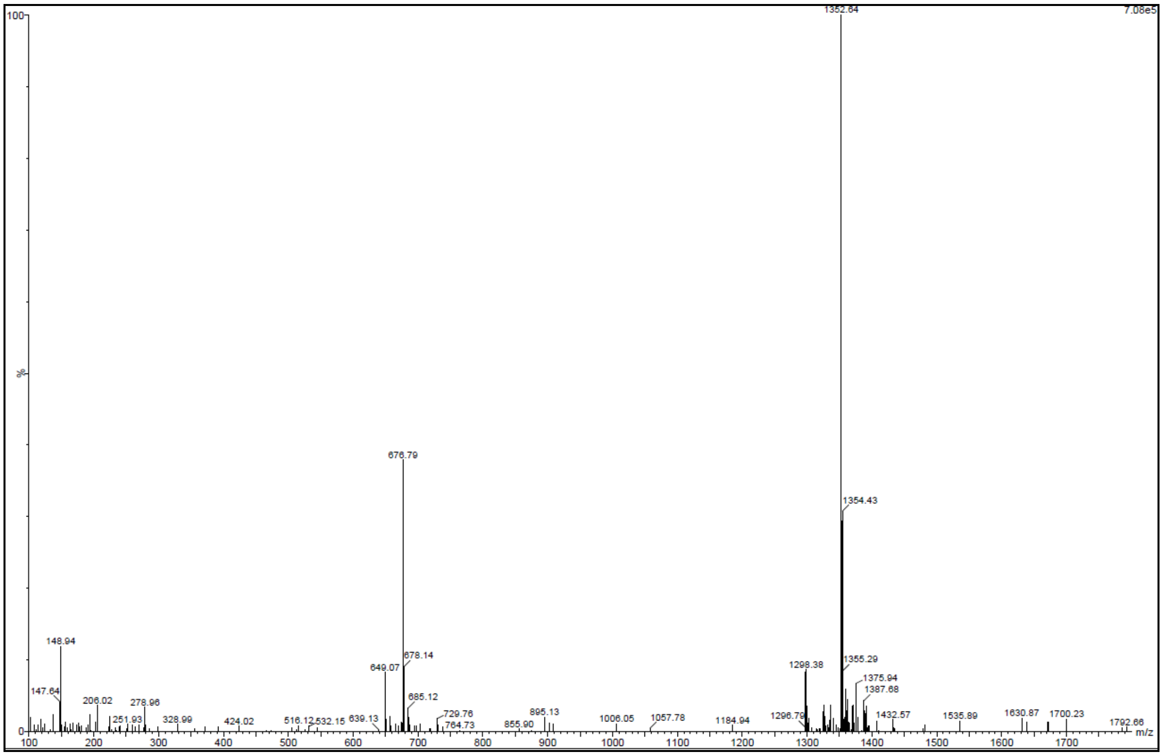


**Figure S8.** Mass spectrum of cCy at 3.72 min elution peak. Expected mass: 1352.42. Observed mass: 1352.64.

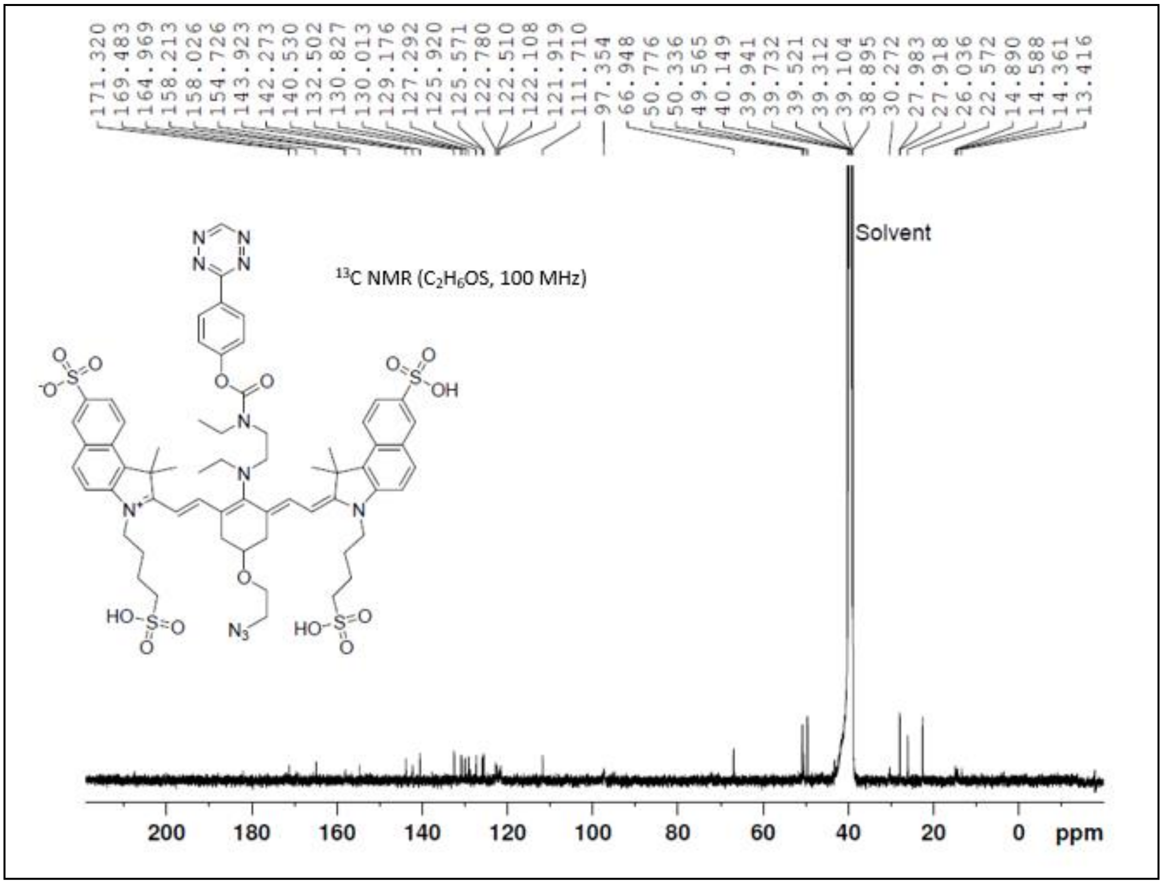


**Figure S9.** $^{13}C$ NMR ($C_2H_6OS$, 100 MHz): δ 13.4, 14.4, 14.6, 14.9, 22.6, 26.0, 27.9, 28.0, 30.3, 49.6, 50.3, 50.8, 66.9, 97.4, 111.7, 121.6, 121.9, 122.1, 122.5, 122.8, 125.6, 125.9, 127.3, 129.2, 130.0, 130.8, 132.5, 140.5, 142.3, 143.9, 154.7, 158.0, 158.2, 165.0, 169.5, 171.3.

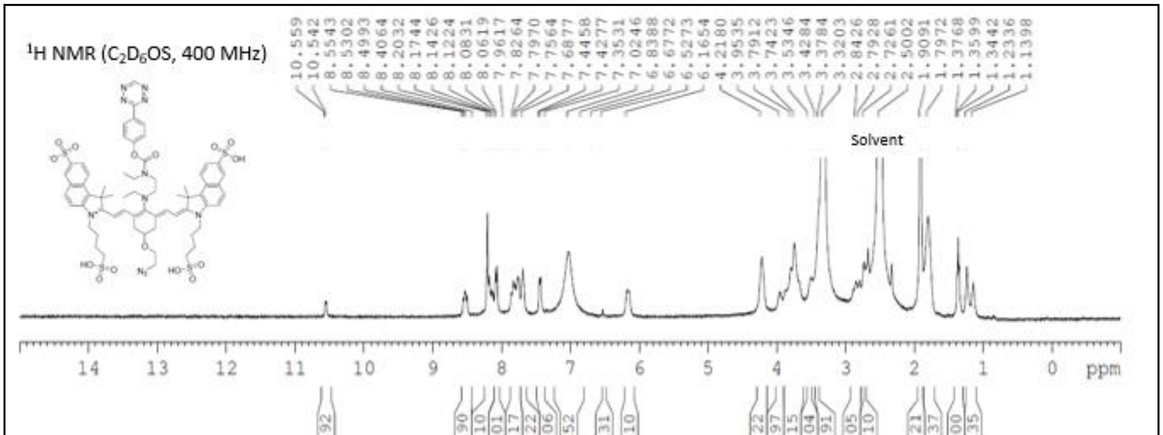


**Figure S10.** $^{1}H$ NMR ($C_2D_6OS$, 400 MHz): δ 1.14 (d,3H, J=36.0 Hz), 1.34 (t, 2H, J= 8.0 Hz), 1.79 (br, 7H), 1.91 (s, 9H), 2.73 (br, 2H), 2.79 (d, 3H, J= 20 Hz), 3.37 (br, 3H), 3.53 (s, 2H), 3.74 (br, 9H), 3.95 (s, 2H), 4.22 (s, 4H), 6.17 (s, 2H), 6.53 (s, 1H), 7.02 (br, 9H), 7.43 (d, 2H, J= 8.0 Hz), 7.69 (s, 2H), 7.76 (t, 4H, J= 16.0 Hz), 8.06 (d, 2H, J=8.0 Hz), 8.12 (q, 3H, J= 8.0 Hz), 8.50 (q, 2H, J= 12Hz).

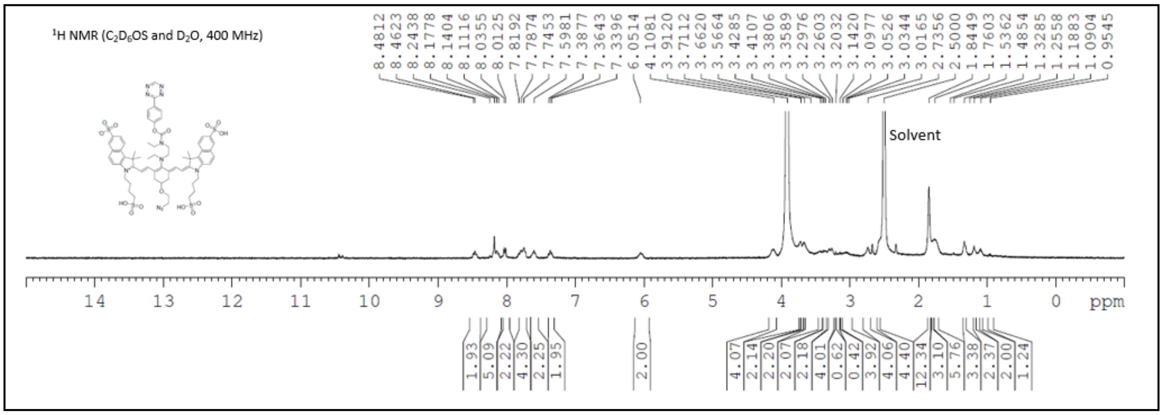


**Figure S11.** $^1$H NMR ($C_2D_6OS$ and $D_2O$, 400 MHz): δ 0.95 (s,1H), 1.09 (s, 2H), 1.19 (s,2H), 1.33 (s, 3H), 1.76 (br, 9H), 1.84 (s, 12H), 2.74 (s, 4H), 3.02 (br, 4H), 3.14 (s, 1H), 3.2 (s, 1H ), 3.26 (d, 4H, J= 16.0 Hz), 3.36 (d, 2H, J= 8.0 Hz), 3.41 (d, 2H, J=8.0 Hz), 3.66 (s, 2H), 3.71 (s, 2H), 4.1 (s, 4H), 6.05 (s, 2H), 7.34 (t, 2H, J= 8.0 Hz), 7.6 (s, 2H), 7.74 (t, 4H, J= 20 Hz), 8.01 (d, 2H, J= 8.0 Hz), 8.04 (br, 5H), 8.46 (m, 2H).
